# University Exams and Psychosocial Stress: Effects on Cortisol Rhythmicity in Students

**DOI:** 10.1101/2021.02.23.432585

**Authors:** Filipy Borghi, Priscila Cristina da Silva, Elisângela Farias-Silva, Fernando Canova, Aglecio Luiz Souza, Aline Barbedo Arouca, Dora Maria Grassi-Kassisse

## Abstract

University students often experience heightened stress during exam periods, which can trigger psychosocial stress and increase cortisol production. This study aims to investigate both the short- and long-term effects of exam-related stress on the hypothalamic-pituitary-adrenal (HPA) axis, focusing on cortisol production and rhythmicity. Twenty-seven undergraduate students (18–24 years) from a biological sciences program participated in this study. Hair cortisol was measured for two months (October and November), while salivary cortisol was collected during final exams in November to assess cortisol rhythmicity. Saliva samples were taken five times per day across three consecutive days. Hair cortisol levels were significantly higher in November, reflecting increased chronic stress during exam periods. However, salivary cortisol maintained a normal circadian rhythm and preserved cortisol awakening response (CAR), despite elevated stress levels. The rhythmicity of cortisol production remained stable across the exam period, though an increase in cortisol before bedtime on the second and third days suggests heightened stress or anticipatory anxiety. Although university exams induce psychosocial stress, students demonstrated resilience in maintaining cortisol rhythmicity and CAR. These findings suggest adaptive stress responses in students, mitigating the risk of stress-related mental health issues. Further research using hair cortisol analysis could provide insights into cumulative stress exposure and aid in developing preventive mental health strategies for university students.

## 1. INTRODUCTION

Stress can be defined as a mismatch between external demands and the perceived ability to cope with the stressor (Gallagher, et al., 2014; Moore, Truscott, St Clair, & Klingborg, 2007). It is largely influenced by individual’s perceptions and cognitive appraisal, making it subjective in nature and difficult to predict objectively (Moore, Truscott, St Clair, & Klingborg, 2007). While stress responses are generally adaptive and play a vital role in enabling individuals to meet external challenges, chronic or overwhelming stress can have detrimental effects (García-León, Pérez-Mármol, Gonzalez-Pérez, García-Ríos, & Peralta-Ramírez, 2019). The neuroendocrine hypothalamus-pituitary-adrenal (HPA) axis is the primary mediator of the body’s response to stress, with cortisol being its main downstream effector (Stalder, et al., 2017).

Systemic cortisol levels are highly variable and are affected by the circadian rhythm, acute stress situations, and pulsatile secretion (Wester & van Rossum, 2015). Traditional methods of measuring cortisol in saliva, blood, or urine only provide snapshots of acute levels over relatively short sampling periods—typically 12 to 24 hours—thus limiting the capacity to capture chronic stress responses (Stalder, et al., 2017; Wester & van Rossum, 2015). Furthermore, these assessments depend on strict adherence to sampling protocols by the patient, which can be challenging. In contrast, hair analysis offers a more stable and reliable biomarker for assessing chronic stress. It has been employed for decades to measure exposure to environmental toxins and drugs and has recently gained attention as a tool for evaluating long-term HPA axis activity (D’Anna-Hernandez, Ross, Natvig, & Laudenslager, 2011; Sauvé, Koren, Walsh, Tokmakejian, & Van Uum, 2007; Wester & van Rossum, 2015). Hair cortisol analysis is particularly advantageous as it reflects endocrine activity over previous months, offering insights into past stress events (Stalder, et al., 2017).

Undergraduate students are frequently exposed to significant stressors throughout their university experience. Research has shown that stressful situations in academic settings are associated with reduced academic performance, lower grade point averages, decreased graduation rates, and higher rates of dropout, alongside heightened psychological distress (Dwyer & Cummings, 2001; Elias, Ping, & Abdullah, 2011; Portoghese, et al., 2019). The pressure to perform well during exam periods is a notable trigger of psychosocial stress, contributing to elevated cortisol release (Karatsoreos & McEwen, 2014; Liston, McEwen, & Casey, 2009). Given the increasing levels of stress among undergraduates, this study aims to explore both the short- and long-term effects of exam periods on the HPA axis in this population by measuring cortisol production over time.

## 2. METHODS

### 2.1 Subjects

Volunteers aged between 18 and 24 years were invited to participate during class and through posters that outlined the general inclusion criteria. They signed a free and informed consent form, declaring their understanding of the procedures that would be performed during the protocol. The study was conducted in accordance with the guidelines established in the Declaration of Helsinki, and ethical approval was granted by the Research Ethics Committee of the School of Medical Sciences/University of Campinas (CAAE: 41225614.8.0000.5404). This study was performed with the collaboration of biological sciences undergraduate students from the University of Campinas (Unicamp). This study involved the collaboration of biological sciences undergraduate students from the University of Campinas (Unicamp). The selection of this undergraduate course was intentional, as both full-time and evening programmes follow the same curriculum, with nearly all subjects taught within the same institute, thereby minimizing the stress associated with commuting within the university. All samples were collected during the final exam/test week at the end of the academic year.

### 2.2 Hair cortisol samples

Twenty-seven undergraduate students from the biological sciences course at Unicamp provided hair samples ranging from 3 and 5 centimetres in length. The hair strands were collected from the posterior vertex of the scalp and cut as close to the scalp as possible, following Manenschijn et al. (Manenschijn, Koper, Lamberts, & van Rossum, 2011). Hair cortisol extraction was performed according to Meyer et al. (Meyer, Novak, Hamel, & Rosenberg, 2014). The results are presented in nmol/L and were derived from 40 mg of hair samples, which were sprayed and resuspended in 0.2 ml of assay buffer, and then converted to pg/mg of hair. It was assumed that the average hair growth rate is 1 cm per month, as reported by Dolnick et al. (Dolnick, Lindahl, Terrill, & Reynolds, 1969). The monthly production of cortisol was evaluated for each volunteer over the preceding two months, October and November.

### 2.3 Salivary cortisol samples

Participants were instructed to collect saliva samples at home using Sallivetes®, which were to be stored under refrigeration. Twenty-eight undergraduate students collected saliva to assess cortisol rhythmicity. Cortisol rhythmicity was evaluated through saliva collections taken at five time points during the day over three consecutive days in November (Tuesday to Thursday): upon awakening (6 a.m.), 30 minutes after awakening (6:30 a.m.), before lunch (noon), before dinner (6 p.m.), and before bedtime (11 p.m.). The samples were centrifuged at 2,800 rpm for 20 minutes at 4°C and analysed using a commercial ELISA kit (Diagnostics Biochem Canada Inc. - Ref CAN-C-290) at the Laboratory of Stress Studies (LABEEST) - UNICAMP, following the procedures of Madeira et al. (Madeira, et al., 2019). The results were expressed in nmol/L for each sample and as the area under the curve for all saliva samples.

### 2.4 Sample size justification and sensitivity analysis

The sample size for this study included 27 undergraduate students providing hair samples and 28 students providing saliva samples. The sample size was determined based on practical constraints, including participant availability and the logistical challenges associated with repeated cortisol measurements. Although an a priori power analysis was not conducted, a post-hoc sensitivity analysis was performed to estimate the smallest detectable effect size given the study’s design and sample size. The analysis indicated that the study is capable of detecting effect sizes of approximately 0.78 (for hair cortisol) and 0.76 (for saliva cortisol) with 80% power at a significance level of α = 0.05. These results suggest that the study is sufficiently powered to detect medium-to-large effects.

### 2.5 Statistical analysis

Data are presented as means ± SEM. Normality was confirmed by the D’Agostino & Pearson test. Paired or unpaired Student’s t-tests were performed for parametric data, and Wilcoxon or Mann-Whitney tests were performed for nonparametric data. All statistical analyses were conducted using GraphPad Prism version 8.00 for Windows (GraphPad Software, San Diego, California, USA). The level of significance was set at p<0.05. Sensitivity analysis was performed using G*Power software to determine the smallest effect size detectable given the study’s sample size.

## 3. RESULTS

### 3.1 Hair cortisol and psychosocial stress

To verify the chronic effect of test/exam weeks on cortisol release, we evaluated hair segments from two different months: one without tests/exams (October) and one with tests/exams and possible emergence of psychosocial stress (November). The results showed that hair cortisol levels in November were significantly higher than those observed in October (Figure 1). This finding indicates a potential cumulative effect of stress during the exam period, reinforcing the notion that psychosocial stress may lead to sustained elevations in cortisol levels.

**Figure 1.**
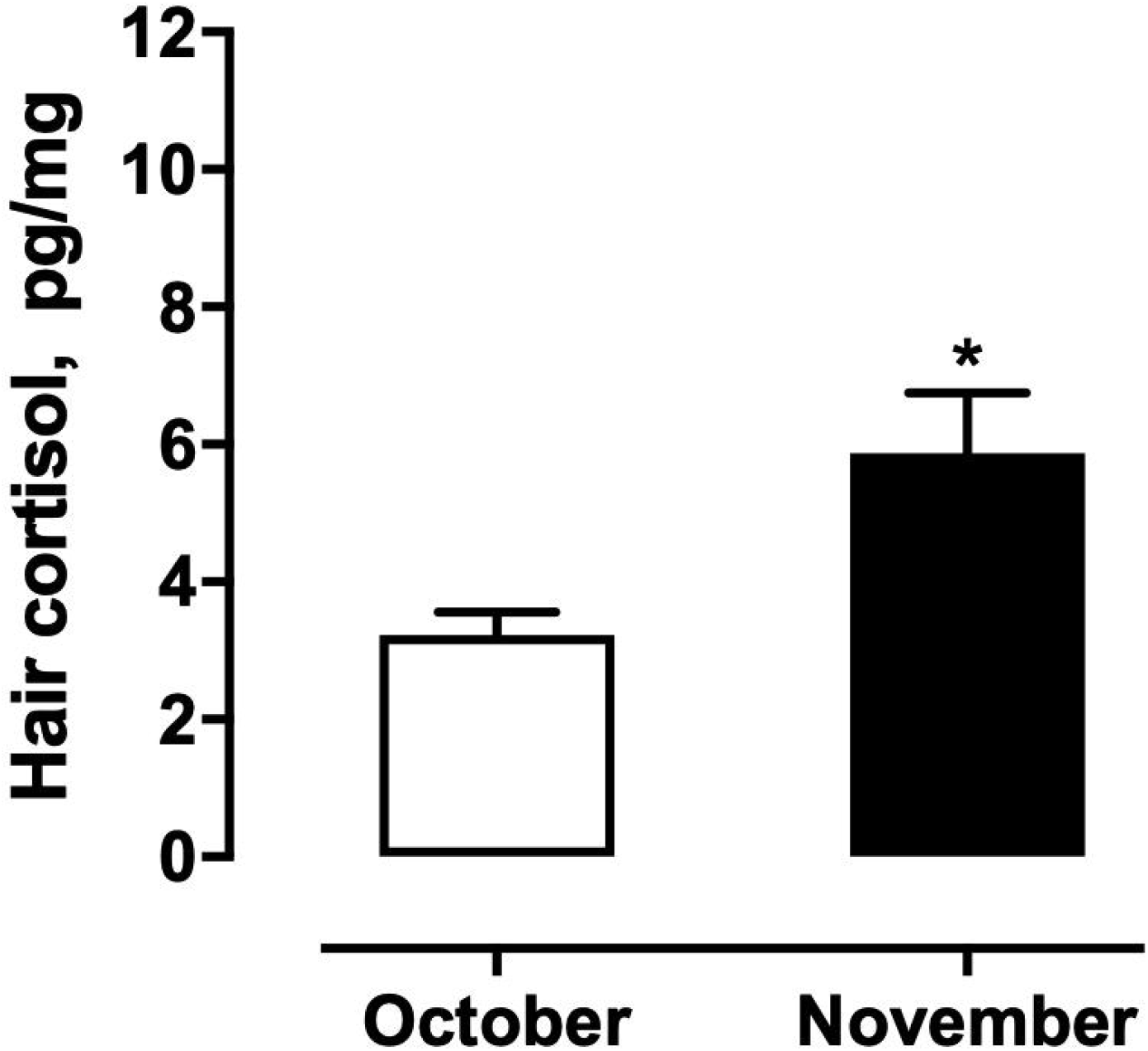
Hair cortisol levels in undergraduate students in October (a month without exams) and November (a month with exams). Data are presented as mean ± SEM. *p < 0.05 within the same group.

### 3.2 Salivary cortisol

These volunteers also showed rhythmicity in cortisol release over the three consecutive sample days in November, the test/exam period, with higher values in the morning and a significant decrease throughout the day. Despite maintaining adequate rhythmicity during the test period, only the 1^st^ day exhibited a significant difference between the before-dinner sample and the before-bedtime sample, suggesting an increase in cortisol on the 2^nd^ and 3^rd^ days at the before-bedtime collection point when compared to the 1^st^ day. This finding can indicate heightened stress or anticipatory anxiety as the test period progressed, potentially reflecting the concept of anticipatory stress or progressive allostatic load. There was no significant change in the amount and rhythmicity of cortisol produced on different days, supporting the notion that while the overall pattern was maintained, specific stress-related elevations occurred at certain times.

The cortisol awakening response (CAR) was preserved among the volunteers, with significantly higher cortisol levels 30 minutes after awakening on all sample days. Although CAR values increased over the three days, the change was not statistically significant. This stability in CAR suggests that the volunteers’ HPA axis reactivity to awakening was not markedly altered by the test/exam stress, indicating resilience in their acute stress response mechanisms, despite the context of increased overall stress during exam weeks (Figure 3).

**Figure 2.**
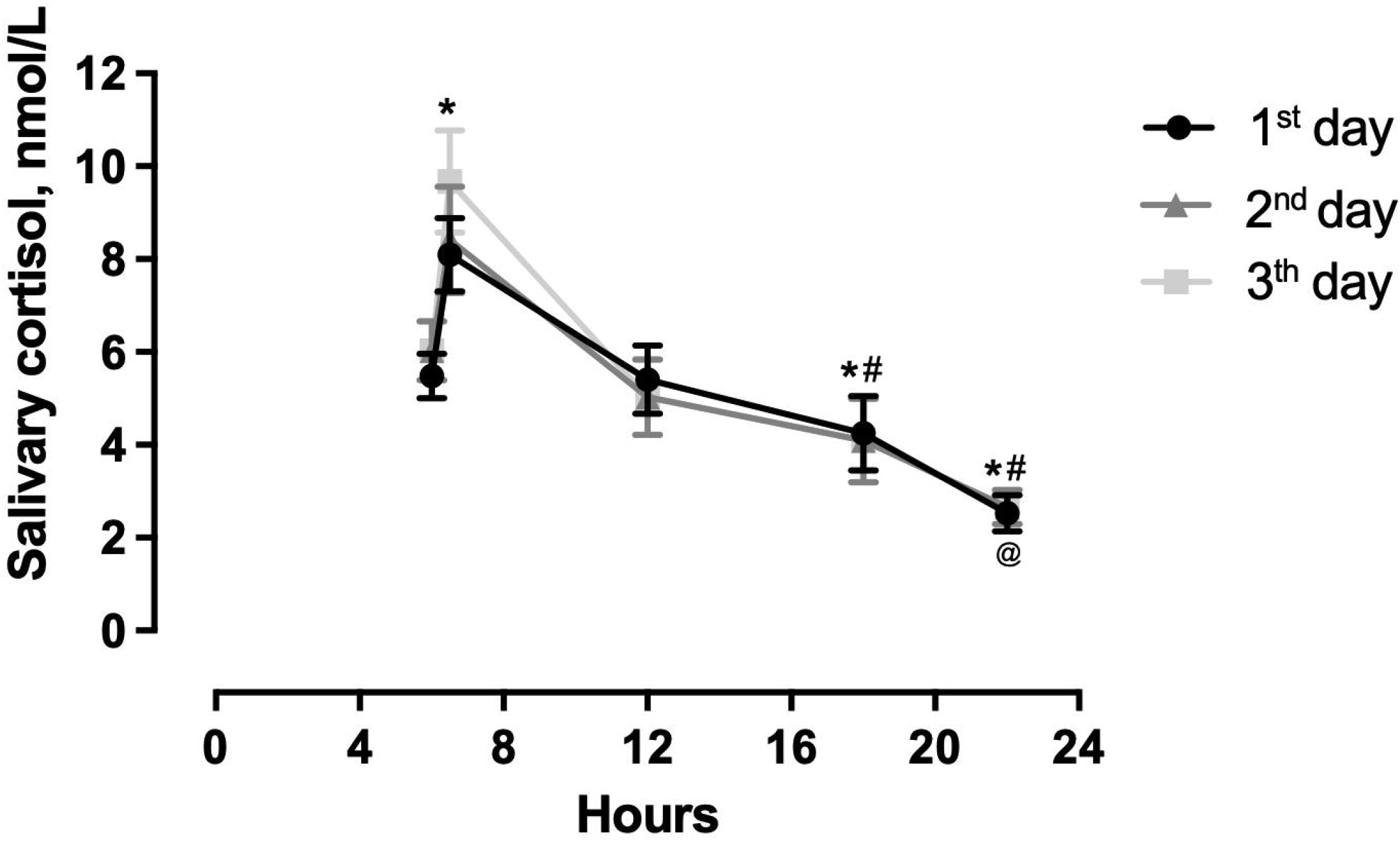
Cortisol rhythmicity from saliva samples collected at five different times of the day over three consecutive days: upon awakening (6 a.m.), 30 minutes after awakening (6:30 a.m.), before lunch (noon), before dinner (6 p.m.), and before bedtime (11 p.m.). Data are presented as mean ± SEM. *p < 0.05 vs. 6 a.m. within the same group; ^#^p < 0.05 vs. noon within the same group; ^@^p < 0.05 for 1^st^ day 6 p.m. vs. 1^st^ day 11 p.m.

**Figure 3.**
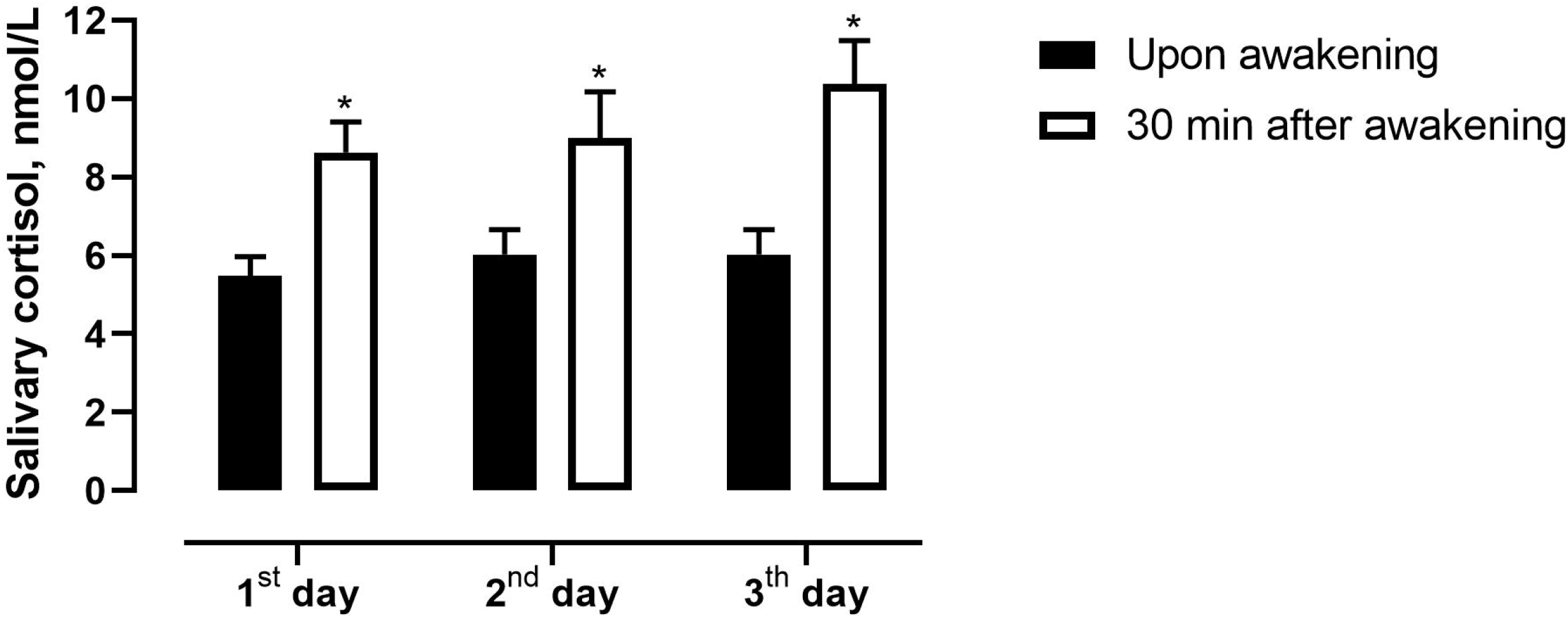
Cortisol Awakening Response (CAR) in undergraduate students over different consecutive days. Data are presented as mean ± SEM. *p < 0.05 within the same group.

## 4. DISCUSSION

Hair cortisol provides a valuable retrospective reflection of cortisol release over several months. In recent years, substantial evidence has accumulated regarding key aspects of hair cortisol measurements, including their validity as indicators of long-term systemic cortisol concentrations, their reliability through repeated assessments, and their robustness against various influences. Based on these findings, researchers have begun using cortisol measurements to address more specific questions about the role of stress in long-term cortisol production and its impact on various health-related conditions. Furthermore, the potential to measure the long-term production of other steroid hormones (e.g., androgens or estrogens) through hair analysis presents an intriguing prospect for future research. Given its unique characteristics, hair analysis holds great promise for significantly enhancing our understanding of steroid hormones (Braig, et al., 2016; Serwinski, Salavecz, Kirschbaum, & Steptoe, 2016; Stalder, et al., 2012; Stalder, et al., 2017).

The analysis of hair cortisol levels provided evidence of the chronic effects of test/exam periods on cortisol release. By comparing hair segments from October (a month without tests/exams) and November (a month with tests/exams and potential psychosocial stress), we observed a significant increase in hair cortisol levels in November. This finding aligns with the understanding that hair cortisol acts as a cumulative measure of HPA axis activity over the preceding months, reflecting prolonged exposure to stress (Wright, Hickman, & Laudenslager, 2015). The elevated levels in November indicate that the academic stress associated with tests/exams has a discernible impact on chronic cortisol production.

A major challenge in hair cortisol research is the significant variation in estimates, even among participants of similar age, in relatively small, non-population-based studies. This issue of “convenience sampling” may limit the generalizability of our results, as it often leads to biases that may influence the outcomes. Future studies could address this limitation by utilizing more representative sampling methods. Despite this variability, the results obtained in this study for hair cortisol align with the literature for a healthy population. For example, a study measuring hair cortisol in Mexican women teachers and women from the Icelandic SAGA pilot cohort found average hair cortisol readings of 6.0 pg/mg and 4.7 pg/mg, respectively (Lynch, et al., 2022). Another study analysing healthy children aged 0-18 years from the Netherlands reported lower values until age 6, with gradually increasing values reaching 3.0 pg/mg by age 18 (de Kruijff, et al., 2020). Additionally, a study using a Brazilian cohort showed hair cortisol concentrations of 7.8 pg/mg for children and 5.6 pg/mg for their mothers (Martins, et al., 2023). Our volunteers showed 3.2 pg/mg in October and 5.8 pg/mg in November. These findings reinforce that hair cortisol can be a reliable indicator of long-term HPA activity, reflecting both chronic stress and the natural variability found in different populations and age groups.

Daily variations in stressors, such as differences in workload between weekdays and weekends, can obscure long-term changes in cortisol rhythms (Adam, Hawkley, Kudielka, & Cacioppo, 2006; Karlamangla, Friedman, Seeman, Stawksi, & Almeida, 2013). Research literature has reported associations between cortisol parameters and psychosocial outcomes in studies using 3-day sampling protocols (Hulett, Fessele, Clayton, & Eaton, 2019). Through this protocol, we observed that our volunteers showed increased evening cortisol over time, which may be linked to fatigue, as suggested by other studies (Schmidt, et al., 2016; Tell, Mathews, & Janusek, 2014; Weinrib, et al., 2010). This increase in evening cortisol could also indicate anticipatory stress as exam periods approach, reflecting a progressive allostatic load on the body.

The HPA axis can be assessed by quantifying salivary cortisol 30 minutes after awakening (Fries, Dettenborn, & Kirschbaum, 2009; Pruessner, et al., 1997; Wang, et al., 2014). Cortisol values 30 minutes after awakening that are significantly higher than those upon awakening indicate that the HPA axis is functioning normally. The CAR mechanism is independent of cortisol secretion throughout the day and is influenced by factors that are not yet fully understood. However importantly, it is abnormalities in cortisol secretion rhythms, whether in CAR or overall rhythmicity, that are associated with various diseases, psychosocial variables, and chronic stress (Clow, Thorn, Evans, & Hucklebridge, 2004). The CAR cortisol values are consistent with those described in the literature for a healthy population, reflecting a 38% increase from baseline values upon awakening, with a healthy response considered up to 100% of this baseline (Dedovic & Ngiam, 2015; Vreeburg, et al., 2010). Therefore, our findings demonstrate that the studied population, despite experiencing psychosocial stress, maintained normal cortisol rhythmicity and a preserved CAR. This stability in CAR suggests resilience in acute stress response mechanisms, indicating that the population may effectively manage short-term stress despite academic pressures. The rhythmicity maintained throughout the week confirms that the studied population is healthy with regard to salivary cortisol evaluation.

Biological instruments and techniques for evaluating chronic stress, such as hair cortisol measurement, are essential for understanding its impact on the body and its association with psychiatric disorders. Given the high incidence of mental illness among undergraduate students (Campos, Oliveira, Mello, & Dantas, 2017), there is a pressing need for accurate biological markers to quantify the effects of psychosocial stress. In comparison to similar studies, cortisol levels in this undergraduate population are comparable to those observed in the Canadian elderly population and lower than those found in healthy third-year U.S. medical students during successive clinical rotations (Souza-Talarico, Plusquellec, Lupien, Fiocco, & Suchecki, 2014; Tseng, Iosif, & Seritan, 2011). Although the absence of standardized international baselines complicates direct comparisons, this pilot study makes a valuable contribution toward establishing such baselines, offering researchers a critical tool for assessing mental health in specific populations. These comparisons suggest that different demographics may exhibit varying levels of resilience or vulnerability to psychosocial stress, highlighting the importance of context in interpreting cortisol data.

Nevertheless, this pilot study has several limitations, including a small sample size, adherence issues throughout the protocol, and the use of a convenience sample, which reduces the generalizability of the results. Despite these challenges, the study’s unique design—focusing on both hair cortisol and salivary rhythmicity during a highly stressful period—provides novel insights into stress biomarkers. While larger sample sizes are recommended for future replication, the sensitivity analysis indicates that the study is adequately powered to detect medium-to-large effect sizes (approximately 0.76 to 0.78). Consequently, the main findings remain relevant and provide a valuable foundation for further research in this field. Future studies should aim to address these limitations by refining the identification of biological instruments and techniques for chronic stress evaluation and its relationship with detrimental effects on the organism. Collecting data on hair and saliva cortisol across different life stages, along with relevant clinical variables, could be instrumental in developing specific cortisol profiles for clinical staging of mental health conditions (Koumantarou Malisiova, et al., 2021). Additionally, future research should explore how these findings can inform mental health interventions at universities, enhancing the practical significance of stress biomarker studies.

## 5. CONCLUSION

Exams week in universities can become a detrimental factor for undergraduate health, as it often results in elevated cortisol levels throughout this period. While this period is a known trigger for psychosocial stress, it did not induce significant short-term changes in cortisol production rhythmicity. This suggests that young people may adapt to such stressors, potentially mitigating their impact on mental health. Additionally, further investigations into this population, particularly regarding their previous experiences of allostatic overload as assessed by hair cortisol levels, could provide valuable insights for developing preventive mental health strategies within the university.

## 6. ACKNOWLEDGEMENT

The authors wish to thank all the volunteers who participated in the study. We thank Tarsilo Onuluk for his thoughtful comments on the manuscript. This study was supported by Coordenação de Aperfeiçoamento de Pessoal de Nível Superior - Brasil (CAPES – Finance Code 001), Serviço de Apoio ao Estudante da Unicamp (SAE-Unicamp), Fundo de Apoio ao Ensino, Pesquisa e Extensão (FAEPEX-PRP), Conselho Nacional de Desenvolvimento Científico e Tecnológico (CNPq), and Fundação de Amparo à Pesquisa do Estado de São Paulo (FAPESP). Parts of these results were presented at the XXIV Congresso de Iniciação Científica da Unicamp (PIBIC) – 2016 and the 17º Congresso de Stress do ISMA-BR – 2017.

